# Immediate targeting of host ribosomes by jumbo phage encoded proteins

**DOI:** 10.1101/2023.02.26.530069

**Authors:** Milan Gerovac, Kotaro Chihara, Laura Wicke, Bettina Böttcher, Rob Lavigne, Jörg Vogel

**Author notes:** Corresponding author: Jörg Vogel.

## Abstract

Bacteriophages must seize control of the host gene expression machinery to promote their own protein synthesis. Since the bacterial hosts are armed with numerous anti-phage defence systems, it is essential that mechanisms of host take-over act immediately upon infection. Although individual proteins that modulate components of the bacterial gene expression apparatus have been described in several different phages, systematic approaches which capture the phage’s arsenal for immediate targeting of host transcription and translation processes have been lacking. In particular, there are no known phage factors that associate directly with host ribosomes to modulate protein synthesis. Here, we take an integrative high-throughput approach to uncover numerous new proteins encoded by the jumbo phage ΦKZ that target the gene expression machinery of the Gram-negative human pathogen *Pseudomonas aeruginosa* immediately upon infection. By integrating biochemical and structural analyses, we identify a conserved phage factor that associates with the large ribosomal subunit by binding the 5S ribosomal RNA. This highly abundant factor is amongst the earliest ΦKZ proteins expressed after infection and stays bound to ribosomes during the entire translation cycle. Our study provides a general strategy to decipher molecular components of phage-mediated host take-over and argues that phage genomes represent a large discovery space for proteins that modulate the host gene expression machinery.

## Introduction

Bacteriophages hold great promise for use in biotechnological applications and antibacterial therapy because they modify fundamental processes of life to subvert their bacterial host. Jumbo phages present particularly exciting opportunities to identify molecular factors crucial to a successful host take-over since they feature extraordinarily large genomes (200-600 kb) that encode hundreds of proteins of currently unknown function. One interesting model is ΦKZ, a jumbo phage that infects *Pseudomonas aeruginosa*, a Gram-negative bacterium and a major cause of nosocomial and antibiotic-resistant infections (Jernigan et al., 2020; Mulani et al., 2019). ΦKZ shares a strikingly complex infection cycle with several other jumbo phages, involving the formation of a pseudo-nucleus that spatially separates genome replication and transcription from translation in the cytosol (Chaikeeratisak et al., 2017). The pseudo-nucleus is said to exclude DNA-targeting host defence systems rendering the phage resistant to certain CRISPR-Cas systems (Malone et al., 2019; Mendoza et al., 2020).

Phages need mechanisms that allow them to immediately seize control of the host gene expression machinery not only to proceed through the infection process efficiently but also to quickly produce factors to antagonize host defence systems. Bacteria encode numerous host defence systems, several of which sense phages immediately after infection (Bernheim & Sorek, 2020). Over the years, bottom-up studies of individual ΦKZ proteins have provided evidence that this phage uses multiple strategies to promote and protect the expression of its own genome, e.g., specialised phage-encoded RNA polymerases (Ceyssens et al., 2014) or an inhibitor of the major host endoribonuclease (Van den Bossche et al., 2016). Still, we have only preliminary knowledge of the true number and temporal activity of such phage-encoded factors, especially those that act in the first line of host take-over. Moreover, phage proteins that directly modify translation by targeting the bacterial ribosome have not been identified to date (De Smet et al., 2017).

Here, we undertook a systematic analysis of cellular protein complexes in ΦKZ-infected *Pseudomonas* cells to identify phage factors that engage the host gene expression machinery early during infection. Strikingly, we observed several proteins that associate with the host polymerase or translating ribosomes. The molecular characterization of one of these ribosome-associated phage factors shows that it acts during early infection and targets a core region of the bacterial protein synthesis machinery during host take-over.

### Phage proteins fractionate with the host gene expression machinery

To systematically identify phage proteins that interact with and subvert the host gene expression machinery, we used Grad-seq (Smirnov et al., 2016). This method predicts molecular complexes based on the separation of cellular lysates by a classical glycerol gradient, followed by high-throughput RNA sequencing (RNA-seq) and mass-spectrometry (MS) analyses of the individual gradient fractions. Our earlier work established that major *P. aeruginosa* protein complexes remain intact upon phage infection (Gerovac et al., 2021). Here, to capture early host take-over events, we infected *P. aeruginosa* with ΦKZ for 10, 20 and 30 minutes and analysed pooled samples in a 10-40% glycerol gradient. This size range captures monomeric proteins below 100 kDa up to 4 MDa ribosomes in twenty fractions plus the pellet. These fractions were then analysed by MS. As previously described, subunits of >200 kDa metabolic complexes, the host RNA polymerase (RNAP) and ribosomes can be identified based on correlated sedimentation profiles (Gerovac et al., 2021) (**Fig. 1a, Ext. Data 1a**). While the apparent bacterial proteome appears largely unaffected by infection when read on standard gels, quantitative MS analysis identified ∼10% of the total protein count as phage-derived (**Ext. Data 1b-d**).

**Fig. 1.**
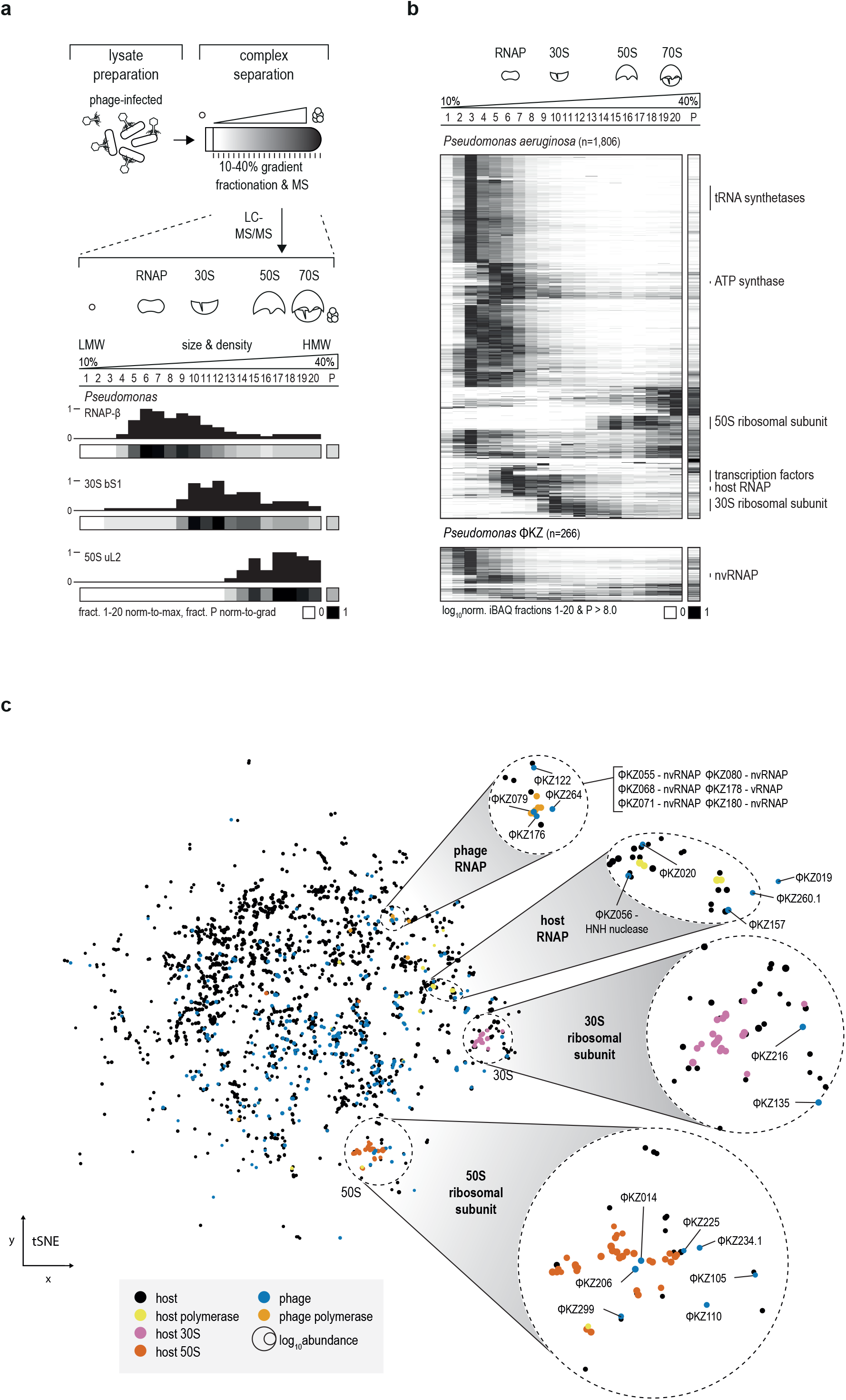
ΦKZ proteins associate with the host gene expression machinery. **a**, *Pseudomonas* cells were infected with ΦKZ. Cellular lysates were analysed in a glycerol gradient by size and density in 20 fractions plus the pellet that cover monomeric proteins, polymerases, up to ribosomes. Mass spectrometric quantification yielded sedimentation profiles of individual proteins, e.g. RNAP β subunit, 30S bS1 and 50S protein uL2. In the colour-coding, fractions 1-20 are normalised to the maximum and the pellet fraction P is normalised to the protein abundance in the gradient. **b**, Host proteins in high-molecular-weight fractions correspond to gene expression machinery subunits. In line, ΦKZ viral polymerase subunits sedimented in fraction five. (rel. log_10_iBAQ>8.0) **c**, Clustering of sedimentation profiles revealed subunits that form large macromolecular complexes. ΦKZ polymerase subunits clustered together with ΦKZ079/122/176, and -264 that were previously not associated to have a role in transcription. In the cluster for the host RNA-polymerase, ΦKZ019/020/56/157, and -260.1 co-sedimented. ΦKZ216 was suggested to co-sediment with the 30S ribosomal subunit. Several ΦKZ factors clustered with proteins from the 50S ribosomal subunit, e.g. ΦKZ014/105/110/206/225/234.1, and -299. In general, phage factors appear to cluster with many diverse complexes.

Our sampling approach enabled us to detect almost three quarters (266 out of 369) of the annotated ΦKZ proteins (at rel. log_10_iBAQ>8, **Suppl. Table 1, Fig. 1b**). Like host proteins, most phage proteins are present in the first three low-molecular-weight (LMW) fractions, suggesting that they are monomeric or form small <100-kDa complexes. Nevertheless, ∼30 phage proteins are present in high-molecular-weight (HMW) fractions (**Fig. 1b**). These include six annotated structural virion proteins that also accumulated in the pellet fraction, indicating that they are likely part of virion assembly intermediates (**Ext. Data 1e**).

A global clustering analysis of protein sedimentation profiles revealed the association of phage proteins with key gene expression complexes of *P. aeruginosa*, i.e., RNAP and the small (30S) and large (50S) ribosomal subunits (**Fig. 1c**). For example, five phage proteins that have so far not been associated with transcription are present in RNAP fractions, suggesting their potential interaction with host RNAP. Many jumbo phages also encode their own RNAPs (McAllister & Raskin, 1993; Sokolova et al., 2020). A virion RNA polymerase (vRNAP) is injected together with the phage genome for initial phage genome transcription, while a phage-encoded non-virion RNA polymerase (nvRNAP) takes over transcription at later stages of infection in a host-RNAP-independent fashion (Ceyssens et al., 2014; Sokolova et al., 2020). We find that ΦKZ RNAP subunits (proteins ΦKZ055, -068, -071, -073, -074, -080, - 123, -178, and -180 (Ceyssens et al., 2014)) cluster together, which further validates our clustering analysis. Proteins ΦKZ079, -122, -176, and -264 are also present in this cluster, suggesting that they are interaction partners of the nvRNAP. Strikingly, we also observed that nine phage proteins peak in ribosomal fractions, but were absent from the pellet (**Fig. 1c, Ext. Data 1e**), making them strong candidates for ribosome-targeting factors.

### Early expressed phage proteins form stable complexes with ribosomes

The nine ΦKZ proteins observed in ribosomal fractions were predicted to target either the 30S (ΦKZ135, -216) or the 50S subunit (ΦKZ014, -105, -110, -206, -225, -234.1, - 299) (**Fig. 1c**), and to the best of our knowledge, are of unknown function. To prioritize candidates that act on host protein synthesis immediately upon infection, we performed a high-resolution gene expression analysis, isolating total RNA of *P. aeruginosa* before and every two minutes after adding ΦKZ, followed by RNA-sequencing. We focussed on the first 10 minutes of the ∼1-h ΦKZ infection cycle, which are crucial to establishing a productive infection. During this time, host DNA is degraded and phage DNA replication is initiated (Chaikeeratisak et al. 2021). A small pseudo-nucleus forms at one of the cell poles. It grows during later replication cycles, when it is centred in the middle of the cell by the phage-derived spindle-apparatus (Erb et al., 2014).

Our RNA-seq time-course revealed that phage mRNAs were expressed remarkably rapidly and accounted for ∼40% of all coding transcripts after 10 minutes (**Fig. 2a**). The most abundant transcripts at that time point included several that encode proteins present in ribosomal fractions, e.g., ΦKZ014, -105, and -216 (**Fig. 2b**). Importantly, these mRNAs accumulate faster than the transcripts encoding nvRNAP subunits, the pseudo-nucleus protein ChmA or the spindle-apparatus protein PhuZ, suggesting ribosome targeting by ΦKZ during early host take-over is a high priority.

**Fig. 2.**
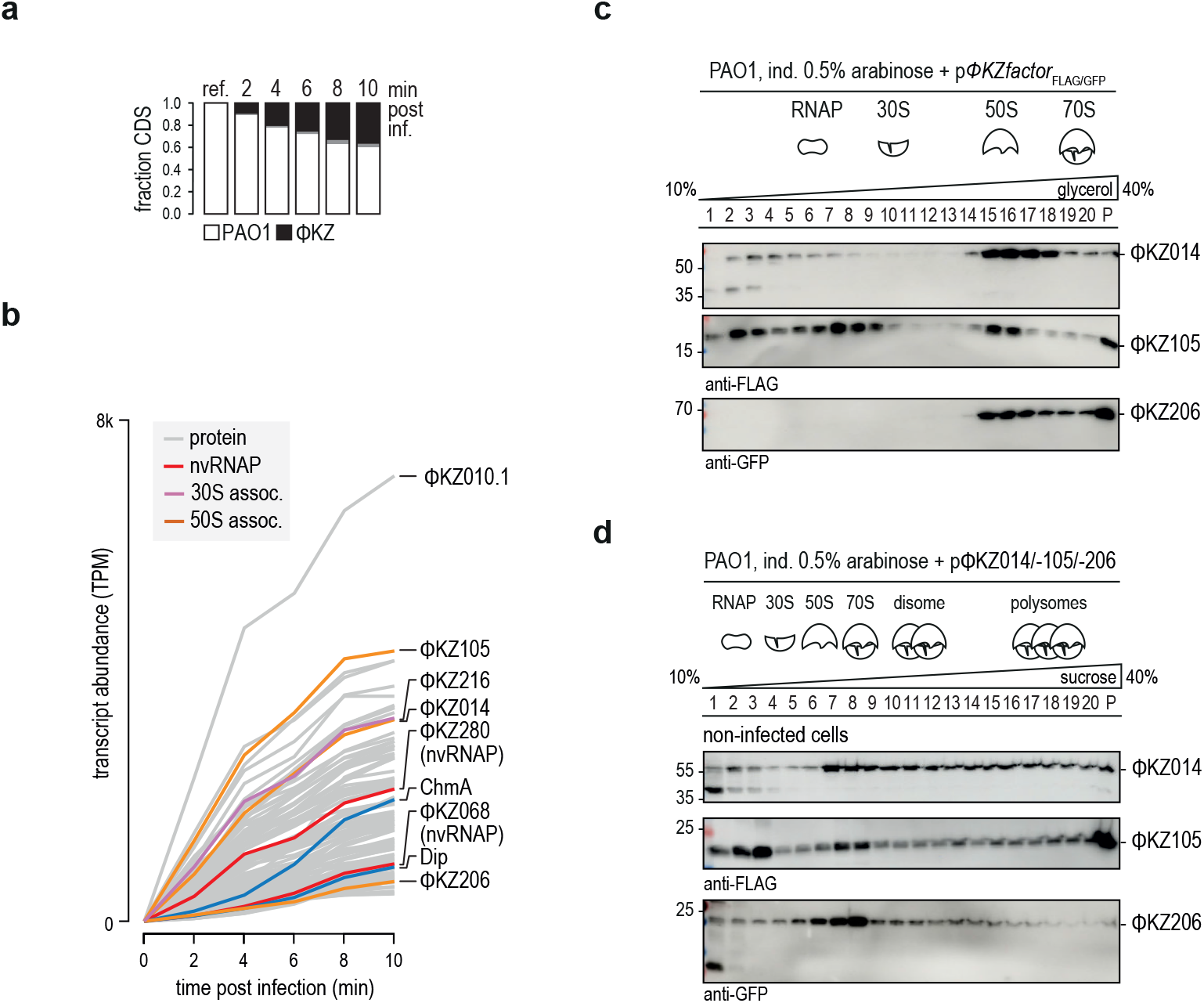
Early expressed phage proteins target translating ribosomes. **a**, In the first ten minutes after ΦKZ infection, phage transcripts accumulate to ∼40%. **b**, Among the most abundant early transcripts, several encode for factors that appeared to sediment in ribosomal fractions (**Fig. 1b,c**). Strikingly, the nvRNAP subunits ΦKZ068/-280 the RNA-degradosome inhibitor Dip and the pseudo-nucleus protein ChmA are less expressed than the ribosome associated factors. Cut-off at sum of most abundant transcripts >85% total phage reads, n=115. **c**, After ectopic expression of ΦKZ014, - 105, and -206 these factors sediment in ribosomal fractions similar to the MS-determined profiles. **d**, In addition, ΦKZ014 and -105 accumulated in polysome fractions indicating a role in translation.

Based on these observations, we selected the proteins ΦKZ014 and ΦKZ105 to probe for their interaction with translating ribosomes. We also included ΦKZ206, a protein that showed a very similar sedimentation profile to ΦKZ014, but whose expression peaked later. To confirm the sedimentation profile of these proteins, we ectopically expressed FLAG-tagged ΦKZ014, -105, and -206 in uninfected *P. aeruginosa* and their position in the glycerol gradient by immunoblotting was analysed (**Fig. 2c**). We observed similar sedimentation profiles as in the Grad-seq experiment, indicating that the association of these proteins with ribosomal fractions is independent of infection and of other phage proteins. In addition, ΦKZ014 and ΦKZ105 accumulated in polysome fractions indicating a role in actively translating ribosomes (**Fig. 2d**).

### ΦKZ014 is an abundant protein that targets the 50S

We focused on ΦKZ014, because the presence of homologs of this protein in many ΦKZ-like phages (Phabio, SL2, ΦPA3, OMKO1, Psa21, EL, KTN4, PA1C, PA7, Fnug, vB_PaeM_PS119XW, PA02, based on PHROG-ID: 16056, PVOG-ID: 9777, **Fig. 3a**) suggested a conserved function. Western blot analysis using a ΦKZ014 antibody and probing *P. aeruginosa* samples from an infection time course detected the protein as early as 5 min post-infection (**Fig. 3b**), in line with the early abundance of the ΦKZ014 mRNA (**Fig. 2b**). Protein expression plateaued after 10 minutes, at which stage the ΦKZ014 protein reached ∼2,000 copies per cell, as inferred from a quantitative comparison of western blot signals for the protein in phage-infected cells with a standard provided by purified recombinant ΦKZ014 protein (**Fig. 3c**).

**Fig. 3.**
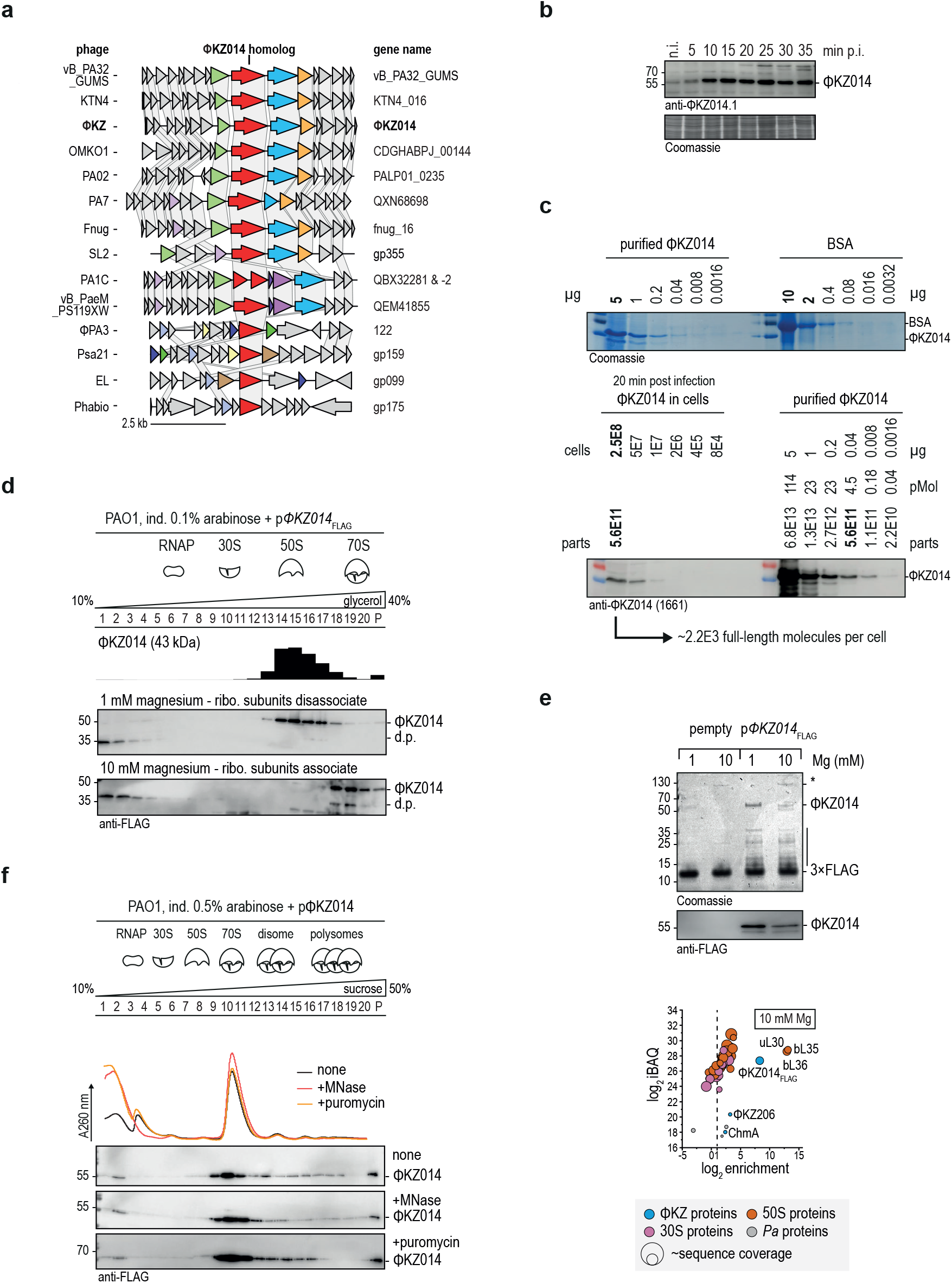
ΦKZ014 is conserved, abundant, and interacts directly with the large ribosomal subunit. **a**, ΦKZ014 appears to be conserved in a number of other ΦKZ-like phages. **b**, ΦKZ014 is 5 min post-infection detectable in the immunoblot and accumulates to saturation at 10 min. **c**, ΦKZ014 reaches 20 min post infection about two thousand copies per cell. The copy number of ΦKZ014 per cell was quantified by quantification of a purified denatured ΦKZ014 protein to bovine serum albumin (BSA) and immunoblot relation to the protein level post infection. **d**, ΦKZ014 sediments in large ribosomal subunit fractions at 1 mM magnesium where ribosomal subunits are dissociated. When subunits remain joined at 10 mM magnesium ΦKZ014 sediments in 70S fractions indicating an interaction with the ribosome. **e**, Pull-down of ectopically expressed ΦKZ014 from infected cells enriched for 50S subunit proteins uL30, bL35, bL36, confirming an interaction with 50S subunit. **f**, ΦKZ014 interaction with the mRNA transcript or the nascent chain was excluded by MNase cleavage of RNA and nascent chain release by puromycylation, both did not cause a shift to top fractions indicating a preserved and direct interaction with the ribosome.

Next, we took two biochemical approaches to narrow down the interaction site of ΦKZ014 on the bacterial ribosome. First, we probed the ribosome interaction of ΦKZ014 at low magnesium ion conditions, when the 70S dissociates into its two subunits. In these settings, ΦKZ014 shifts to the 50S fractions (**Fig. 3d**). At higher magnesium levels, which preserve 70S ribosomes, ΦKZ014 co-migrates with the 70S fractions. This confirms that ΦKZ014 targets the 50S subunit and that the interaction likely occurs outside of the intersubunit site. Secondly, in an orthogonal protein pull-down assay using FLAG-tagged ΦKZ014 followed by MS, ΦKZ014 strongly enriched 50S ribosomal proteins, especially uL30, bL35, and bL36 (**Fig. 3e**).

Next, we probed if the ribosome association of ΦKZ014 was mediated by interactions with mRNA, in which case it would be sensitive to treatment with micrococcal nuclease (MNase), or by the nascent protein chain, in which case it would be sensitive to release by the antibiotic puromycin. When transcripts were degraded by MNase, we observed the expected shift of the phage protein from the polysome fraction to 70S ribosomes, and there was no shift to LMW fractions (**Fig. 3f**). The sedimentation of ΦKZ014 was not altered upon puromycin treatment. Together, these observations confirm a stable and direct association of ΦKZ014 with host ribosomes independent of the transcript or nascent chain.

### The ΦKZ014 protein clamps the terminal end of 5S rRNA

To gain insights into the architecture of the ΦKZ014–ribosome complex, we exogenously expressed HIS-tagged ΦKZ014 in *Pseudomonas* and affinity purified ribosomes via the tagged protein (**Fig. 4a**). The complex was reconstructed by single-particle cryo-EM (**Fig. 4b, Ext. Data 2, Ext. Data Table 1**). Interestingly, we observed that ΦKZ014 occupied the 50S ribosomal subunit at the central protuberance (**Fig. 4b,c**), a region that facilitates communication between various functional ribosome sites (Mohan et al., 2014). The main constituent of the central protuberance is the 5S rRNA, which is essential for efficient translation (Ammons et al., 1999; Huang et al., 2020). Notably, ΦKZ014 occupies the ribosome in different translation states with tRNAs in the A- or E-sites (**Ext. Data 2b**).

**Fig. 4.**
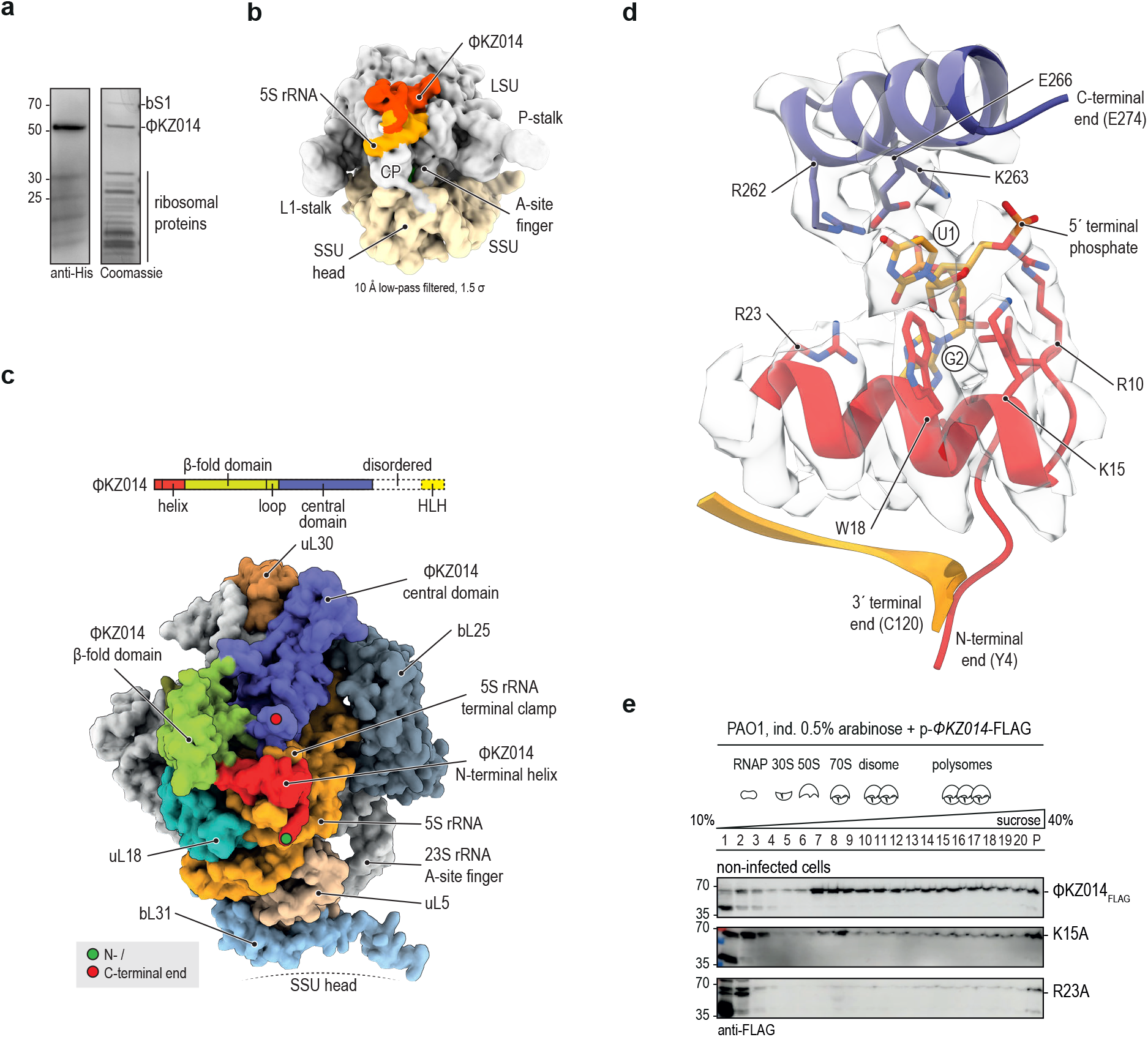
ΦKZ014 clamps the 5S rRNA. **a**, The 70S-ΦKZ014 complex was affinity purified after ectopic expression of ΦKZ014 in *Pseudomonas*. **b**, ΦKZ014 occupies the large ribosomal subunit on top of the 5S rRNA. 10 Å low-pass filtered, 1.5 σ **c**, The N-terminal helix of ΦKZ014 and the last helix of the central domain clamp the terminal end of the 5S rRNA. Both domains are connected by the β-fold domain. The C-terminal end is unstructured and not resolved in the map. **d**, Terminal U1 of 5S rRNA is coordinated by R10, K15, W18 and R262, K263, and E266. This strong coordination flips U1 outside of the helical configuration. The second residue G2 (base-pairing with C118) is coordinated by R23 at the base. 3′-terminal end of 5S rRNA and the N-terminal end of ΦKZ014 are displayed as cartoons without density. **e**, Upon exchange of key Interacting residues from the structural model the interaction with the ribosome was diminished and the proteins accumulated in LMW fractions in a free form. This was especially evident for the Arg23Ala and Lys15Ala exchanges. ΦKZ014 wt reference from Fig. 2d.

ΦKZ014 consists of a N-terminal helix that is connected to a β-fold domain, followed by a central helical domain. Its predicted C-terminal helix-loop-helix domain is linked by a helix and is not resolved in the electron density map (**Fig. 4c, Ext. Data 3a**). Strikingly, ΦKZ014 clamps the terminal end of 5S rRNA and interacts with it via charge-based interactions (**Fig. 4d, Ext. Data 3b**). The first uridine residue of the 5S rRNA is coordinated by the N-terminal ΦKZ014 residues Arg10, Lys15, Trp18 together with the central domain residues Arg262 and Lys263. The second guanosine residue of 5S rRNA is coordinated by Arg23. This multivalent coordination leads to a flip of the uridine out of its helical conformation. ΦKZ014 is also in contact with the 23S rRNA, including the stem of the A-site finger helix (ASF), which is important for the accommodation of tRNA in the A-site. Two sequential histidines within a loop in the central domain (His178 and His179) infiltrate the 23S rRNA major groove at positions 842 and 908 (**Ext. Data 3c**). This direct contact might lead to structural rearrangements in the ASF through rotational extension.

To validate the observed interaction sites, we mutated two coordinating residues of ΦKZ014 to alanine, i.e., lysine 15 and arginine 23. These coordinating residues are well conserved among various ΦKZ014 homologs (**Ext. Data 3d**). Either of these two mutations (K15A, R23A) abrogated the association of the ΦKZ014 protein with ribosomes, causing it to accumulate in LMW fractions (**Fig. 4e**). Thus, our structural data validate the direct interaction of ΦKZ014 with the large ribosomal subunit and suggest that ΦKZ014 might play a role in translation via modulation of 5S rRNA.

### Outlook

Phages and bacteria engage in an arms race that drives diversification and evolution of novel host defence and phage takeover systems (Stern & Sorek, 2011). Jumbo phages whose large genomes typically encode hundreds of functionally uncharacterized and evolutionary untraceable proteins represent a large untapped reservoir of novel non-conserved protein families. However, the sheer number of these uncharacterized proteins requires experimental selection prior to functional analysis. The screening approach introduced here should be applicable to diverse phages in the quest to systematically identify phage proteins that engage with the host gene expression machinery. Coupling this analysis to gene expression profiling early during infection enabled us to select proteins that are likely to be important for immediate host-takeover. A more detailed investigation of these candidates promises to yield new insights into how the phage manipulates its host early during the infection process. To support further exploration of our data sets, we developed an interactive online explorer that allows correlation between host and phage protein sedimentation profiles and enables straightforward predictions of protein interactions (www.helmholtz-hiri.de/en/datasets/gradseqphage).

While phages often provide their own RNAPs to ensure transcription of their genome in the host, for protein synthesis they fully rely on the host translation machinery. This machinery comprises large ribosomal RNAs and >50 protein components, and its size and complex assembly process makes it very unlikely that a phage would provide its own ribosomes, either physically or genetically. Thus, for a phage to seize control of protein synthesis, it must be able to modulate host ribosomes. Indeed, the first prediction that phage proteins might associate with ribosomal subunits dates back more than 50 years (Smith & Haselkorn, 1969; Kutter et al., 2018), but the identity of such phage factors has remained elusive.

Here, we report several ΦKZ proteins that interact directly with the ribosome, as shown exemplary by ectopic expression of three of these proteins in uninfected *P. aeruginosa* cells (**Fig. 2c,d**) and a high-resolution structural analysis of the ΦKZ014–ribosome complex (**Fig. 4**). In contrast to phage-encoded proteins with sequence similarity to known ribosomal or translation factors that were predicted in metagenomic studies (Al-Shayeb et al., 2020; Mizuno et al., 2019) or in phage lambda (Jaskunas et al. 1975), these ΦKZ proteins have no sequence similarity to any known host protein. Some of them, including ΦKZ014, are expressed immediately after infection and are highly abundant, suggesting crucial functions in early host take-over. Such functions might not be restricted to prioritising phage protein production, but might also play a role in accelerating production of factors for neutralisation of host defence systems (Pons et al., 2023). Moreover, given that in ΦKZ transcription and translation are spatially separated by the pseudo-nucleus, one may also envision a coupling of both processes upon translocation of the phage mRNA to the cytosol. Notably, ΦKZ014 plateaus at ∼2,000 copies/cell, fewer than the number of cellular ribosomes (Bremer & Dennis, 2008). Hence, only a fraction of ribosomes will be bound by ΦKZ014 and it is tempting to speculate that ΦKZ014 might localise a subset of ribosomes to the vicinity of the pseudo-nucleus or modify these ribosomes for efficient phage mRNA translation.

The discovery of phage proteins that modify host RNAP has been instrumental in understanding individual steps of the bacterial transcription cycle (Tabib-Salazar et al. 2019). We envision a similar potential for ribosome-targeting phage proteins. For example, our structural data reveal that ΦKZ014 interacts with the 50S subunit in close contact with the 5S rRNA; no other proteins have been observed to interact at this site. As such, understanding how ΦKZ014 reorganises protein synthesis in the first minutes of infection may also help to unveil the molecular function of the universal 5S rRNA. Of note, more than half a century after the discovery of 5S rRNA (Dinman & Dontsova, 2005) and despite its established importance for efficient translation (Ammons et al., 1999) and during ribosome biogenesis (Huang et al., 2020), we still lack a clear mechanistic understanding of how 5S rRNA contributes to protein synthesis. More generally, investigation of the mechanisms of ribosome-targeting phage proteins might also lead to novel biotechnological or medical uses, e.g., better *in vitro* translation systems or inhibitors to recall phages used in phage therapy.

## Methods

### Phage lysate preparation

Bacterial cells were grown to an OD_600_ ∼0.3 at 37 °C in LB media. Cells were infected with phage lysate (high titer 1-3E11 PFU) at an MOI between 0.1-1. In the following three to four hours lysis was observed and the solution was cleared at 5,000 × *g* for 15 min at 4 °C. Phages were precipitated with one volume precipitation buffer (2.5 M NaCl, 20% (w/v) PEG8000) for 1 h at 4 °C, and pelleted at 4,000 *g* for 10 min at 4 °C. Precipitation buffer was carefully removed and the pellet was resuspended in 0.5 volume phage storage buffer (50 mM Tris/HCl pH 7.5, 100 mM NaCl, 16 mM MgCl_2_). Lysates were stored at 4 °C and the phage titre was regularly validated by a plaque assay on PAO1.

### Phage plaque assay

Bacterial cells were grown to OD_600_ ∼0.3 at 37 °C. Soft agar (0.5% agar, 1 mM MgCl_2_) was boiled and allowed to cool down to 42 °C. 100 µl bacterial cells were added to 10 ml soft agar, poured into a petri dish plate, and solidified with a closed lid. The phage lysate was serially diluted in LB media from E-2 to E-9. 5 µl of each phage lysate dilution was spotted onto the soft-agar plate. Overnight plaques became visible that were counted to determine plaque forming units (PFU) per ml phage lysate.

### Glycerol gradient fractionation

*P. aeruginosa* PAO1 was grown to an OD600 ∼0.3 at 37 °C and was infected with ΦKZ at an MOI of 15, as previously described (Gerovac et al., 2021). Phage infection was allowed for 5 min without shaking and then cells were harvested 10, 20, and 30 min post infection on ice. Cells were pelleted and the different time points were pooled to extend our view on complexes in the first half of the replication cycle. 30-90 OD Cells were resuspended in 800 µl lysis buffer (20 mM Tris/HCl pH 7.4, 150 mM KCl, 2.5 mM MgCl_2_, 1 mM dithiothreitol (DTT), 0.1% (v/v) Triton X100, 1 mM phenylmethylsulfonyl fluoride (PMSF)). Cells were lysed in 2 ml FastPrep tubes with lysing matrix E (MP Biomedicals) for 2× 40 s FastPrep-24 instrument (MP Biomedicals) at 6 m/s at 4 °C. The lysate was cleared at 13k×*g* for 10 min and layered onto a 10-40% glycerol gradient in lysis buffer without PMSF. The gradient was centrifuged for 17 h at 100k×*g* and 4 °C. 20 590 µl fractions were collected and the pellet was resuspended extensively in the last one. Samples were mixed with 0.25 volume 4x Bolt™ SDS sample buffer (Invitrogen) and were boiled for 5 min at 95 °C. 2 pmol human albumin was spiked-in per 50 µl fraction for subsequent normalisation. MS sample preparation and measurement was conducted at the Proteomics Core Facility EMBL Heidelberg by Jennifer Schwarz. Samples were reduced and alkylated with 10 mM dithiothreitol (DTT) at 56 °C for 30 min, and 2-chloroacetamide at room temperature in the dark for 30 min, respectively. Samples were cleaned-up using the SP3 protocol (Hughes et al., 2019). 300 ng sequencing grade Trypsin (Promega) in 50 mM ammonium bicarbonate was added for overnight digestion at 37 °C. Peptides were recovered by collecting the supernatant on a magnet followed by a second elution with ultrapure water. Samples were dried under vacuum centrifugation and reconstituted in 10 μl 1% formic acid, 4% acetonitrile and then stored at -80 °C until LC-MS analysis.

### MS analysis

Samples were analysed at the EMBL Proteomics Core Facility (Heidelberg). An UltiMate 3000 RSLC nano LC system (Dionex) fitted with a trapping cartridge (µ-Precolumn C18 PepMap 100, 5 µm, 300 µm i.d. × 5 mm, 100 Å) and an analytical column (nanoEase™ M/Z HSS T3 column 75 µm × 250 mm C18, 1.8 µm, 100 Å, Waters) was coupled directly to a QExactive Plus (Thermo Scientific) mass spectrometer using the Nanospray Flex™ ion source in positive ion mode. Trapping was carried out with a constant flow of 0.05% trifluoroacetic acid at 30 µl/min onto the trapping column for 4 min. Subsequently, peptides were eluted via the analytical column with a constant flow of 0.3 µl/min with increasing percentage of solvent B (0.1% formic acid in acetonitrile) from 2% to 4% in 4 min, from 4% to 8% in 2 min, then 8% to 25% for a further 89 min, and finally from 25% to 40% in another 17 min and from 40% to 80% in 3 min. The peptides were introduced into the Qexactive plus via a Pico-Tip Emitter 360 µm OD × 20 µm ID; 10 µm tip (MSWIL) and an applied spray voltage of 2.2 kV. The capillary temperature was set at 275 °C. Full mass scans were acquired with mass range 350-1,400 m/z in profile mode with resolution of 70,000. The filling time was set at maximum of 20 ms with a limitation of 3×10^6^ ions. Data dependent acquisition (DDA) was performed with the resolution of the Orbitrap set to 17,500, with a fill time of 50 ms and a limitation of 1×10^5^ ions. A normalized collision energy of 26 was applied. Loop count 20. Isolation window 1.7 *m/z*. Dynamic exclusion time of 30 s was used. The peptide match algorithm was set to ‘preferred’ and charge exclusion ‘unassigned’, charge states 1, 5 – 8 were excluded. MS^2^ data was acquired in centroid mode. The raw mass spectrometry data was processed with MaxQuant (v1.6.17.0) (Cox & Mann, 2008) and searched against the database UP000002438, UP000002098 for *Pseudomonas aeruginosa* and ΦKZ phage, respectively. As an internal standard the entry P02768 (albumin of *homo sapiens*) was used in each experiment. Common contaminants were included in each search. Decoy mode was set to revert. Carbamidomethyl I was set as fixed modification, acetylation of N-termini and oxidation of methionine were set as variable modifications. The mass error tolerance for the full scan MS spectra was set to 20 ppm and for the MS/MS spectra to 0.5 Da. A maximum of two missed cleavages was permitted. For protein identification, a minimum of one unique peptide with a peptide length of at least seven amino acids and a false discovery rate (FDR) below 0.01 were required on the peptide and protein level. Match between runs was enabled with standard settings. Quantification was performed using intensities and iBAQ values (Schwanhäusser et al., 2011) calculated as the sum of the intensities of the identified peptides and divided by the number of observable peptides of a protein. Intensities were normalised to the spike-in and the gradient fractions to obtain sedimentation profiles. t-distributed stochastic neighbor embedding (t-SNE) was conducted in Orange (Demsar et al., 2013).

### Sequencing of RNA

PAO1 was grown to OD_600_ 0.3 in LB media. Cells were washed and inoculated 1:3 in M9 minimal media and grown to OD_600_ 0.5. Cells were infected with 15 MOI ΦKZ phage in SM buffer. Cells were lysed with 1% SDS. The lysate was acidified with 0.3 M NaOAc pH 5.2 and RNA was extracted with PCI, chloroform, and precipitated with Ethanol/0.3 M NaOAc pH 6.5. RNA was DNase digested. A 100 ng RNA sample was used for rRNA depletion with RiboCop rRNA Depletion Kit for Mixed Bacterial Samples (META, Lexogen). RNA quality was assessed on a 2100 Bioanalyzer with the RNA 6000 pico kit (Agilent Technologies). Libraries were prepared with CORALL RNA-Seq Library Prep Kit (Lexogen). PCR amplification was conducted for 14 cycles. Libraries were pooled and spiked with 1% PhiX control library. Sequencing was performed at 5 million reads/sample in single-end mode with 150 nt read length on the NextSeq 500 platform (Illumina) using a Mid output sequencing kit. FASTQ files were demultiplexes with bcl2fastq2 v2.20.0.422 (Illumina). Reads were trimmed with curadapt (1.15, (Martin, 2011)). Reads were mapped to genomics sequences NC_002516 (PAO1) and NC_004629 (ΦKZ) and quantified with READemption 0.4.3 (Förstner et al., 2014).

### Expression of phage proteins

PAO1 was transformed as previously described (Choi et al., 2006) with expression plasmids for arabinose-inducible C-terminally -TEV-3×FLAG tagged phage proteins in the backbone of pJN105. Briefly an overnight culture of PAO1 was inoculated from overnight culture to OD_600_ 0.05 in LB media (with 50 µg/ml gentamicin), induced with 0.1-1% arabinose, and grown to OD_600_ 0.3. Optionally, cells were infected with high titre ΦKZ phage (MOI of 15; 1-3E11 PFU/ml), incubated for 5 min at room-temperature, and then at 37 °C at 220 rpm for 5-20 min. The infection was stopped by rapid cool-down on ice. For Ribo-seq experiments cells were pelleted at room-temperature and the pellet was flash-frozen in liquid nitrogen. Cells were pelleted at 6×k*g* for 15 min. The pellet was frozen at -20 °C.

### Purification of ΦKZ014 protein and absolute quantification

ΦKZ014 purification under native conditions resulted in strong contamination with nucleic acids. Application of stringent conditions resulted in loss of the protein. For absolute quantification, which did not require folded protein, protein was purified under denaturing conditions for normalisation in Western blot detection. H6-3C-PHIKZ014 was expressed from pMiG-41 in PAO1. Cells were grown to an OD 0.6 in LB with 50 μg/ml gentamicin at 37 °C and expression was induced with 0.5% arabinose for 2 h. Cells were collected and lysed by sonication for 2×2 min with (50% amplitude, Sonopuls HD 3200, VS 70T tip, Bandelin) in lysis buffer (Tris/HCl pH 8.0, 0.5 M KCl, 2 mM MgCl_2_, 5 mM DTT, 1 mM PMSF). The lysate was cleared at 100×k*g* for 1 h in Ti70 rotor (Beckman Coulter). The pellet was subsequently resuspended in lysis buffer with 8 M Urea and 20 mM imidazole at room-temperature for 1 h. Not solubilised protein was removed by a second pelleting at 100×k*g* for 1 h in Ti70 rotor (Beckman Coulter). The solubilized and denatured protein was loaded on a 5 ml HisTrap HP column (Cytiva), washed with 6 M urea in lysis buffer and eluted with 350 mM imidazole in 6 M urea lysis buffer. ΦKZ014 was concentrated by centrifugal filtration (3 kDa cut-off, Amicon Ultra, Merck-Millipore). Protein concentration of the purified ΦKZ014 was estimated by correlation of band thickness in Coomassie staining with BSA at ∼50 kDa in SDS-PAGE. Native ΦKZ014 in infected cells was detected by Western blot (1:10k anti-ΦKZ014, Eurogentec, #1661; 1:10k anti-rabbit-HRP, Thermo Scientific, #31460) and cross-correlated to the signal of the purified protein. The cell number used as input for the Western was determined by counting colonies and resulted in 2E8 colony forming units (CFU) per ml at OD600∼0.5, 1/40 of a 50 ml culture was loaded per lane, hence 2.5E8 cells. The number of ribosomes per cell was determined by lysis of a defined number of cells, MNase treatment, and gradient fractionation, followed by estimation of 70S ribosomes by the extinction coefficient 3.84E7 M^−1^cm^−1^ that resulted in 4,000 ribosomes per cell at OD600 ∼ 0.3.

### Polysome profile fractionation

10-30 OD Cells were resuspended on ice in 800 µl polysome lysis buffer (20 mM Tris/HCl pH 7.4, 150 mM KCl, 10-70 mM MgCl_2_, 2 mM DTT, 1 mM PMSF, 100 µg/ml lysozyme), and 500 µl 0.1 mM glass beads were added. Lysis was performed in the Retch M200 mill at 30 Hz for 5 min at 4 °C. The supernatant was cleared at 13×k*g* for 10 min and layered onto a 10-50% (w/v) sucrose gradient in polysome lysis buffer without PMSF and lysozyme. The gradient was centrifuged for 16 h at 70,500×*g* and 4 °C. The A260 nm polysome profiles was recorded on a BioComp fractionator. Twenty fractions (590 µl each) were collected and the pellet was resuspended in the last one and taken completely.

### SDS-PAGE

Samples were resuspended in SDS-PAGE loading dye (60 mM Tris/HCl pH 6.8, 0.2 g/ml sodium dodecyl sulfate (SDS), 0.1 mg/ml bromophenol blue, 77 mg/ml DTT, 10% (v/v) glycerol), boiled and stored at 4 °C. Proteins were resolved on 12-15% acrylamide (37.5:1) gels in a discontinuous Laemmli buffer system constituting of the separation gel (375 mM Tris/HCl pH 8.8, 0.1% (w/v) SDS) and a stacking gel (125 mM Tris/HCl pH 6.8, 0.1% (w/v) SDS) which were polymerized with 0.1% (w/v) ammonium persulfate (APS) and 0.001% (v/v) *N,N,N′,N′*-tetramethylethane-1,2-diamine (TEMED), and resolved in the running buffer (25 mM Tris/HCl, 192 mM glycine, 0.1% (w/v) SDS).

### Immunoblotting

SDS-PAGE gels were semi-dry blotted onto methanol-preactivated polyvinylidene (PVDF) membranes for 1 h at 34 mA/gel (20×8 cm) in transfer buffer (200 mM Tris base, 150 mM glycine, 0.1% (w/v) SDS, 20% (v/v) methanol). Membranes were blocked with 5% BSA in TBS-T buffer (20 mM Tris/HCl pH 7.4, 150 mM NaCl, 0.05% Tween20). The following antibodies were used: anti-FLAG (mouse, 1:3k, Sigma, F1804), anti-His (mouse, 1:3k, Sigma, A7058), anti-c-Myc (mouse, 1:1,000, Sigma, M4439), anti-GFP (mouse, 1:1k, Sigma, 11814460001 (Roche)), anti-mouse-HRP (goat, 1:10k, Thermo Scientific, 31430), anti-rabbit-HRP (goat, 1:10k, Thermo Scientific, 31460), anti-ΦKZ014.1 (1660, rabbit, 1:10k, Eurogentec, produced against peptide EQYGESDDTSDESSY, **Ext. Data 3d**) anti-ΦKZ014.2 (1661, rabbit, 1:10k, Eurogentec, produced against peptide TEYDRNHGWNIREKH, **Ext. Data 3d**).

### Comparative gene cluster analysis

Gene homologs of ΦKZ014 were identified by BLAST (NCBI), JackHmmer (EMBL-EBI), and Colab Fold Alpha Fold multiple-sequence-alignment analysis. From indicated genes the loci were obtained from NCBI with additional 3 kb flanking regions. Sequences were submitted to the online comparative gene cluster analysis (https://cagecat.bioinformatics.nl/) that uses cLinker for cluster generation (Gilchrist and Chooi 2021).

### Cryo-EM sample preparation

H6-3C-PHIKZ014 was expressed from pMiG-41 in PAO1. Cells were grown to an OD 0.6 in LB with 50 μg/ml gentamicin at 37 °C and expression was induced with 0.5% arabinose for 2 h. Cells were collected and lysed by sonication for 2×2 min with (50% amplitude, Sonopuls HD 3200, VS 70T tip, Bandelin) in lysis buffer (20 mM Tris/HCl pH 8.0, 500 mM KCl, 16 mM MgCl_2_, 4 mM 2ME, 1 mM PMSF, 20 mM imidazole). The lysate was cleared at 15×k*g* for 20 min. Equilibrated Protino Ni-IDA beads (Macherey-Nagel) were added and rotated for 30 min. Beads were washed three times with 40 bead volumes lysis buffer. Complexes were eluted with 300 mM imidazole and rebuffered in grid buffer (20 mM HEPES/KOH pH 7.5, 150 mM KCl, 16 mM MgCl_2_, 1 mM DTT, 0.1% TritonX100). Quantifoil grids (holey carbon R3/3 with a 2 nm carbon support) were glow-discharge treated at 10E-1 Torr for 45 s. Samples were vitrified using a Vitrobot Mark IV (FEI), 3.5 µl was applied, and after 45 s of incubation at 4 °C and 100% humidity, excess sample was blotted for 6 s and the grids were plunged into liquid ethane at ∼-160 °C.

### Cryo-EM imaging and reconstruction

Electron Cryo Microscopy was carried out in the cryo EM-facility of the Julius-Maximilians University Würzburg. 8,431 micrographs were collected on a Titan-Krios G3 with an X-FEG source and a Falcon III camera with direct electron detection at 300 kV, a magnification of 75k×, a defocus range -1.0 to - 1.8 µm with 2.21 e^−^/fraction and 40 fractions. The magnified pixel size was 1.0635 Å/pixel. All frames were aligned and summed using MotionCor2 (Zheng et al., 2017). Processing was performed with CryoSPARC v3.3.2 (Punjani et al., 2017) (**Ext. Data 2b**). Final maps were auto-sharpened in Phenix (Liebschner et al., 2019) for subsequent model building and structural refinement.

### Model building

*Pseudomonas* 70S ribosome model was prepared by rigid-body docking of the reference model ribosomal subunits (PDB: 6SPG (Halfon et al., 2019)), L31 was removed and homology modelled with Modeller 10.4 (Webb & Sali, 2016) based on *E. coli* reference from PDB: 7N1P (Rundlet et al., 2021). tRNA was also incorporated from PDB: 7N1P. 5S rRNA was modelled and refined in Coot 0.9.8.1 (Casañal et al., 2020; Emsley et al., 2010), together with ΦKZ014 starting from AlphaFold2 and ColabFold (Jumper et al., 2021; Mirdita et al., 2022) and Phenix 1.20.1-4487 (Liebschner et al., 2019) map-to-model predictions. The final model was real-space refined and validated in Phenix. Structures and maps were displayed with UCSF ChimeraX (1.5, (Pettersen et al., 2021)).

## Data availability

MS data are deposited at the ProteomeXchange consortium via the PRIDE partner repository (Perez-Riverol et al., 2022) with the data set identifier PXD038771 for ΦKZ infected cells, respectively. Raw data after MaxQuant and sequencing analysis are listed in **Suppl. Table 1**. Sedimentation data are browsable in a user-friendly browser at www.helmholtz-hiri.de/en/datasets/gradseqphage. Sequencing raw data and coverage files are accessible at Gene Expression Omnibus (Barrett et al., 2012) with the accession number GSE223979, the analysed data are listed in **Suppl. Table 2**. Cryo-EM density maps of 70S–tRNA(P)–ΦKZ014, 70S– ΦKZ014 (focused), 70S–tRNA(E)–ΦKZ014 were deposited at EMDB under accession number EMD-16566. The final model of 70S–tRNA(P)–ΦKZ014 was deposited at RCSB-PDB 8CD1. Strains, oligonucleotides, plasmids, antibodies, software are listed in **Suppl. Table 3**.

## Acknowledgements

We thank Laura Vogel for technical assistance and Anke Sparmann for discussions and for editing the manuscript. Nikolaos Famelis for cloning of plasmids. Andreas Schlosser and Stephanie Lamer from the technology platform mass spectrometry (Würzburg University) and Jennifer Schwarz from the EMBL Proteomics Core Facility (Heidelberg) for helpful discussions and MS. Electron Cryo Microscopy was carried out in the cryo EM-facility of the Julius-Maximilians-University Würzburg (DFG INST 93/903-1). We thank Cihan Makbul, Jochen Kuper, and Christian Kraft from the cryo-EM facility (University Würzburg) for helpful discussions. We thank the Core Unit SysMed at the University of Würzburg for excellent technical support and RNA sequencing. L.W. holds a predoctoral scholarship from FWO-fundamental research (11D8920N). The manuscript was supported by funding from the European Research Council (ERC) under the European Union’s Horizon 2020 research and innovation program (grant agreement No. 819800) awarded to R.L.. The work was funded by Deutsche Forschungsgemeinschaft (DFG) project 465133664 in the Priority Programme “New Concepts in Prokaryotic Virus-host Interactions – From Single Cells to Microbial Communities” (SPP 2330) awarded to J.V..

## Extended Data

**Extended Data 1.**
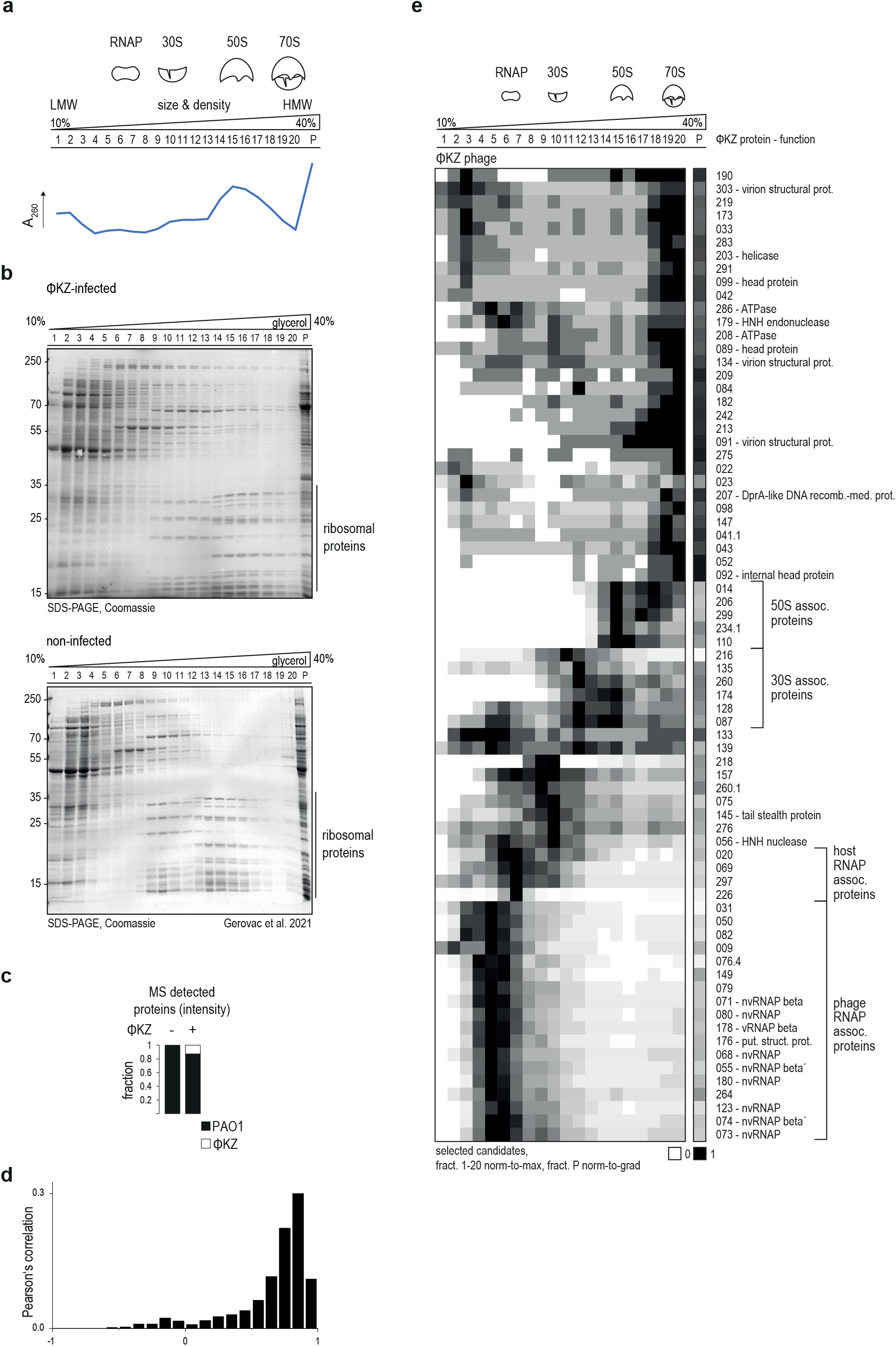
Integrity of the cellular proteome after infection is preserved. **a**, Peaks in the absorption profile of the cellular lysate indicate sedimentation positions of RNAP and ribosomes. **b**, The apparent protein pattern by size is not significantly changed upon infection (non-infected sample, (Gerovac et al., 2021)). **c**, Phage proteins account for ∼10% of total proteins revealed by relative mass-spectrometric quantification by iBAQ. **d**, ∼70% of proteins are >70% correlated in sedimentation compared between infected and non-infected samples (Gerovac et al. 2021). **e**, Sedimentation profiles of selected candidates that appeared in HMW fractions. Phage viral polymerase subunits cluster together in fraction five with several other factors that are not functionally associated with the RNAP. Four factors co-sediment in fraction seven that is known to be dominated by host RNAP. A couple of factors sedimented in fractions associated to ribosomal subunits. Many more factors appear in 70S fractions, but also in the pellet making them likely assembly intermediates of the phage.

**Extended Data 2.**
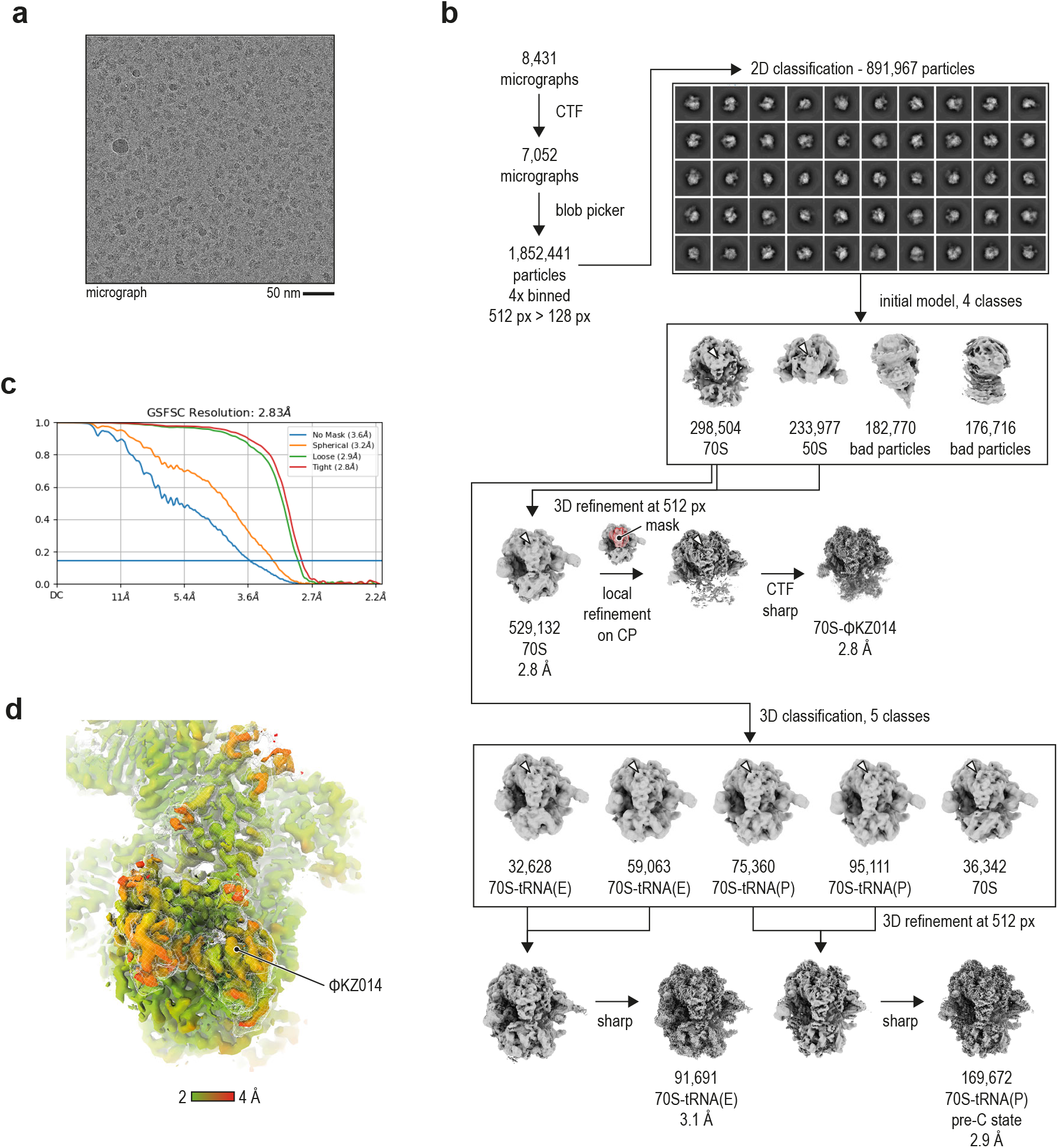
ΦKZ014 occupies translating ribosomes in different states. **a**, Micrograph of the purified 70S-ΦKZ014 complex. **b**, In processing, we selected for classes where ΦKZ014 was associated with the large ribosomal subunit and performed a focused refinement (red mask). Additional classes included the 70S ribosome with tRNA in the P and E site. **c**, FSC curve. The resolution of the focused map was ∼2.8 Å, determined in CryoSPARC refinement. **d**, Local resolution estimation of the additional density that was attributed to ΦKZ014, determined in CryoSPARC.

**Extended Data 3.**
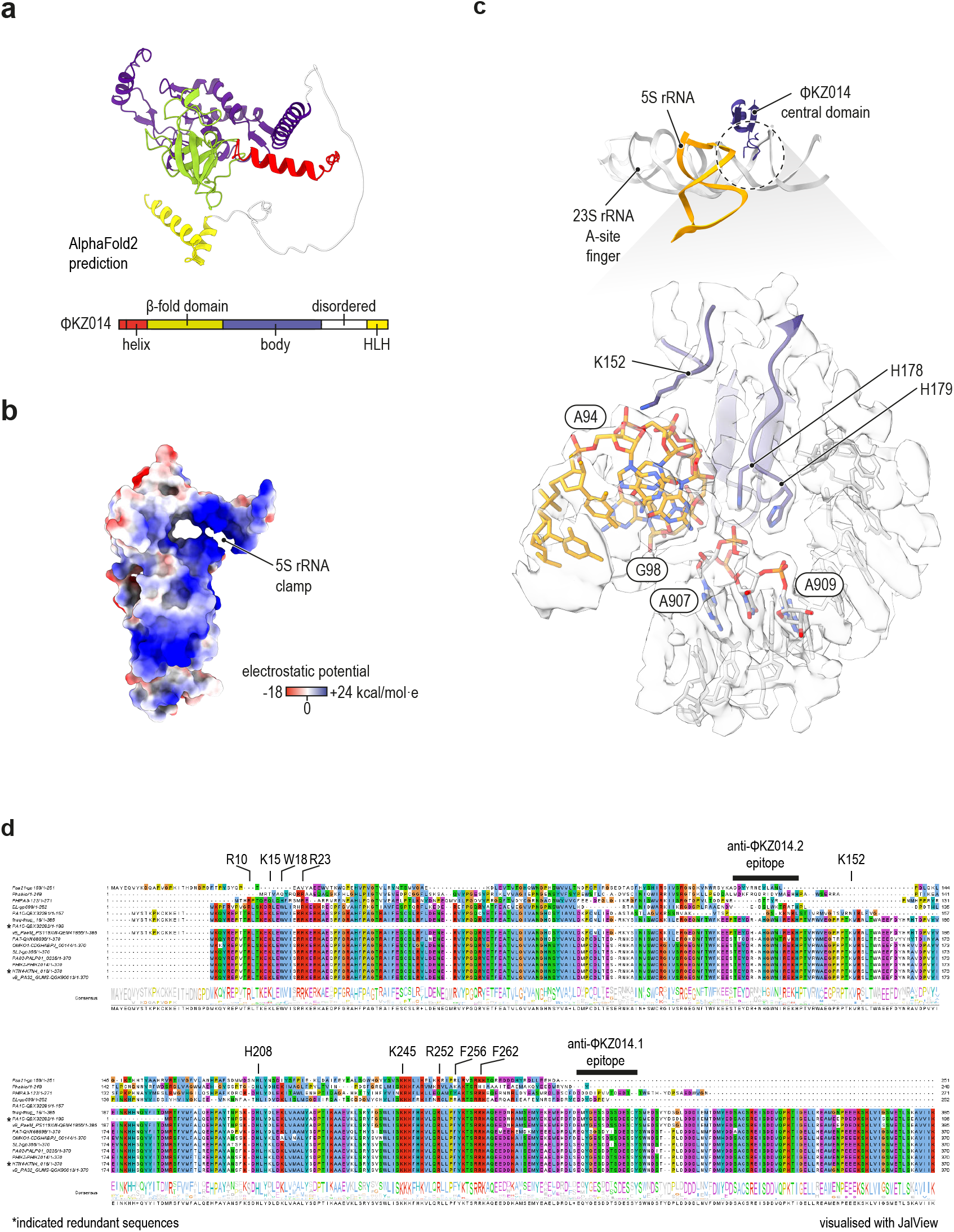
ΦKZ014 structure and interaction sites. **a**, The predicted structure of ΦKZ014 suggested an N-terminal helix, a β-fold domain, a central domain, a disordered region and a C-terminal helix-loop-helix (HLH) domain, determined with AlphaFold2/ColabFold. **b**, The interaction between ΦKZ014 and 5S rRNA is charge-based evident by electrostatic potential of the ΦKZ014 interface that fits to the negatively charged backbone of 5S rRNA. **c**, ΦKZ014 interacts with 23S rRNA backbone at positions 842 and 908 via the two sequential His178 and His179 residues. This is in close proximity to 5S rRNA A94-G98. Lys152 points here in the direction of 5S rRNA backbone. **d**, The key interacting residues are apparently conserved in ΦKZ014 homologs. Interestingly, homologs from phages Psa21, Phabio, EL, and ΦPA3 are more diverse in sequence and the homolog in the phage PA1C appears to be split into two consecutively encoded genes.

**Extended Data Table 1.** Cryo-EM statistics.

|  | #1 70S-tRNA(P)-ΦKZ014 | #2 70S-ΦKZ014 (focused) | #3 70S-tRNA(E)-ΦKZ014 |
| --- | --- | --- | --- |
| EMDB code | EMD-16566 | ----- | ----- |
| PDB code | 8CD1 |  |  |
| **Data collection and Processing** |  |  |  |
| Magnification | 75 k× | 75 k× | 75 k× |
| Voltage (keV) | 300 | 300 | 300 |
| Electron exposure (e<sup>-</sup>/Å<sup>2</sup>) | 19.35 | 19.35 | 19.35 |
| Defocus range (μm) | -1.0 to -1.8 | -1.0 to -1.8 | -1.0 to -1.8 |
| Pixel size (Å) | 1.0635 | 1.0635 | 1.0635 |
| Symmetry imposed | C1 | C1 | C1 |
| Micrographs collected | 8,431 | 8,431 | 8,431 |
| Micrographs used | 7,052 | 7,052 | 7,052 |
| Initial particle images (no.) | 1,852,441 | 1,852,441 | 1,852,441 |
| Final particle images (no.) | 169,672 | 529,132 | 91,323 |
| Map resolution (Å) | 3.0 | 2.8 | 3.1 |
| FSC threshold | 0.143 | 0.143 | 0.143 |
| Map resolution range (Å) | 3.0-3.7 | 2.8-3.6 | 3.1-3.8 |
| **Refinement** in PHENIX 1.20.1-4487 |  |  |  |
| Initial model used (PDB code) | 6SPG |  |  |
| CC mask | 0.80 |  |  |
| Model resolution (Å) | 3.1 |  |  |
| FSC threshold | 0.5 |  |  |
| Map sharpening B factor (Å<sup>2</sup>) | 116.9 |  |  |
| Model composition |  |  |  |
| Chains | 53 |  |  |
| Non-hydrogen atoms | 144,801 |  |  |
| Protein residues | 5,897 |  |  |
| Nucleotides | 4,605 |  |  |
| Ligands | 171 |  |  |
| B factors (Å<sup>2</sup>) (min/max/mean) |  |  |  |
| Protein | 0.19 / 640.79 / 75.69 |  |  |
| RNA | 0.00 / 284.87 / 80.97 |  |  |
| Ligand | 0.00 / 136.39 / 35.38 |  |  |
| R.m.s. deviations |  |  |  |
| Bond lengths (Å) (n) | 0.003 |  |  |
| Bond angles (°) (n) | 0.695 |  |  |
| Validation |  |  |  |
| MolProbity score | 1.66 |  |  |
| Clashscore | 3.16 |  |  |
| Poor rotamers (%) | 1.65 |  |  |
| CaBLAM outliers (%) | 3.97 |  |  |
| EMRinger Score (ΦKZ014<sup>4-274</sup>) | 3.39 |  |  |
| Ramachandran plot |  |  |  |
| Favored (%) | 94.30 |  |  |
| Allowed (%) | 5.61 |  |  |
| Disallowed (%) | 0.09 |  |  |

## Supplementary Material

**Supplementary Table 1 ∣ Grad-seq sedimentation profiles Supplementary**

**Table 2 ∣ Early phage transcripts in ΦKZ-infected cells Supplementary**

**Table 3 ∣ Materials**

